# Blood pressure and the kidney cortex transcriptome response to high sodium diet challenge in female nonhuman primates

**DOI:** 10.1101/2021.10.29.466291

**Authors:** Angelica M. Riojas, Kimberly D. Spradling-Reeves, Robert E. Shade, Sobha R. Puppala, Clinton L. Christensen, Shifra Birnbaum, Jeremy P. Glenn, Cun Li, Hossam Shaltout, Shannan Hall-Ursone, Laura A. Cox

**Author notes:** **Correspondence:** Laura A. Cox, PhD, E-mail address, Address: Wake Forest School of Medicine, Medical Center Boulevard, Winston-Salem, NC 27107, Fax: NA. **Authors’ contributions** RS, KR, LC, and SHU setup the animal study portion related to telemetry collection of baboon BP and kidney biopsy collections. CC, SB, and JG, assisted with BP and kidney biopsy collections and processed kidney biopsies to for sequencing. HS helped with BP analysis and interpretation. KR, LC, and SP contributed to interpretation of BP and transcriptome related results. CL performed IHC and imaging of kidney samples. AR designed study, analyzed and interpreted BP and transcriptome data, performed ELISAs, and wrote the manuscript. All authors read and contributed to the approved final manuscript.

## Abstract

Blood pressure (BP) is influenced by genetic variation and sodium intake. Ninety percent of Americans consume more than the AHA recommended amount of sodium. Studies on the impact of genetic variation and sodium intake on BP in nonhuman primates (NHP) to date have focused on males. *We hypothesized that variation in renal transcriptional networks correlate with BP response to high dietary sodium in female baboons*. Sodium-naïve female baboons (n=7) were fed a low-sodium (LS) diet for 6 weeks followed by a high sodium (HS) diet for 6 weeks. Sodium intake, serum 17 beta-estradiol, and ultrasound-guided kidney biopsies for RNA-Seq BP were collected at the end of each diet. BP was continuously measured for 64-hour periods throughout the study by implantable telemetry devices. On the LS diet, Na^+^ intake and serum 17 beta-estradiol concentration correlated with BP.

Kidney transcriptomes differed by diet; analysis by unbiased weighted gene co-expression network analysis revealed modules of genes correlated with BP on the HS diet. Cell type composition of renal biopsies was consistent among all animals for both diets. Network analysis of module genes showed causal networks linking hormone receptors, proliferation and differentiation, methylation, hypoxia, insulin and lipid regulation, and inflammation as regulators underlying variation in BP on the HS diet. Our results show variation in BP correlated with novel kidney gene networks with master regulators *PPARG* and *MYC* in female baboons on a HS diet. Identification of mechanisms underlying regulators that influence BP will inform better therapies towards greater precision medicine for women.

## Introduction

Sodium intake is known to influence blood pressure (BP) and hypertension (HT) risk^1,2^. Previous studies have identified genetic variants that contribute to BP regulation; however, these variants do not account for all genetic variation regulating BP response to sodium^3–6^. Although some variants have been identified, there are still many people who do not respond to current therapies, thus there is a need for identification of additional functional variants regulating BP^1,7^. Much of what we know about HT is derived from studies that utilized rodent models; however, some genetic mechanisms regulating BP in rodents differ from humans^5,8–10^. For example, gene expression differences between primates and rodent kidneys occur in early development and are maintained in adulthood^11^. Sodium is known to impact the circadian rhythm of BP cycles - nocturnal rodents do not present the characteristic BP drop at nighttime seen in diurnal primates^12,13^. NHP share genetic and physiological characteristics with humans and are appropriate to identify primate-specific genetic variation that influences BP response to sodium^13–20^. In this study, we use sodium-naïve primates, which are almost impossible to find in a human population, to better understand BP regulation with low and high sodium diets and to identify molecular changes that precede HT. These findings are relevant to understanding mechanisms underlying early blood pressure increases that precede HT in humans^13–20^.

Pedigreed, phenotyped and genotyped baboons (*Papio hamadryas*) at the Southwest National Primate Research Center (SNPRC) have been used to study complex heritable human diseases, including HT and sodium-sensitive HT^13–20^. Previous studies of sodium sensitive HT in baboons were limited to males, despite the disease impacting both sexes^1,15,18,21,22^. It was recently established that sex differences exist in sodium naïve baboon kidneys, further supporting a study of high sodium impact on female BP^23^. This sexual dimorphism is observed among many species, and HT models including sodium sensitive HT^24^. Despite these differences, there are no clinical guidelines supporting female specific treatment plans for HT^1^. The goal of this study was to address this knowledge gap by identifying transcriptional networks and genes regulating these networks that correlate with BP response to sodium in female baboons. *We hypothesized that variation in renal transcriptional networks correlate with BP response to dietary sodium in female baboons*.

To investigate this, we challenged pedigreed, genotyped female sodium naïve baboons (*Papio hamadryas*) with a high sodium diet and quantified BP and renal transcriptomic responses. Sodium naïve female baboons maintained on a low sodium (LS) diet throughout their lives were fed a high sodium (HS) diet challenge for 6 weeks. BP was measured continuously for 64-hours by implantable telemetry device for 6 weeks prior to, and for 6 weeks during, a high sodium challenge. Kidney cortex biopsies, blood, and urine samples were collected at the end of the LS phase and HS diet challenge. Blood and urine chemistries were performed, along with transcriptome profiling of kidney cortex biopsies. Sodium intake, BP, and serum 17 beta-estradiol were significantly correlated with each other on the LS diet, but not the HS diet. Conversely, the kidney cortex transcriptome also differed by diet, and unbiased Weighted Gene Co-expression Network Analysis (WGCNA) identified genes associated with BP variation for the HS diet, but not on the LS diet. We identified genes related to hormone receptors, proliferation and differentiation, methylation, hypoxia, insulin and lipid regulation, and inflammation as regulators of the kidney cortex transcriptome correlated with BP on the HS diet. We validated transcript networks findings by immunohistochemistry (IHC) and observed VEGFA in the cortex correlated with blood pressure, sodium intake, and serum 17 beta-estradiol concentration on a high sodium diet.

This study demonstrates significant variation in BP and kidney transcriptome among female baboons fed a HS diet. The similarity between baboons and humans is important for development of diagnostics and therapies that address the role of sodium in BP regulation in women. In addition, our study reveals putative regulatory networks underlying early BP changes with exposure to HS diet.

## Methods

### Ethics, study design, and data collection

The olive baboons utilized in this study (*Papio hamadryas*; Taxonomy ID 9557) were maintained as part of the pedigreed baboon colony at SNPRC, at the Texas Biomedical Research Institute (TBRI), San Antonio, Texas. The TBRI’s Institutional Animal Care and Use Committee (IACUC) reviewed and approved all animal procedures. The SNPRC facilities at the TBRI and animal use programs are accredited by Association for Assessment and Accreditation of Laboratory Animal Care International (AAALAC), which operate according to all National Institutes of Health (NIH) and U.S. Department of Agriculture (USDA) guidelines and are directed by veterinarians (DVM). SNPRC veterinarians made all animal care decisions and enrichment was provided on a daily basis by the SNPRC veterinary staff and behavioral staff in accordance with AAALAC, NIH, and USDA guidelines. Premenopausal female baboons were selected based on family history of HT as previously described (n=7, age 17.86 ± 1.35 years)^18,23^. The KINSHIP program from Pedigree Database System (PEDSYS v. 2.0) was used with the Stevens-Boyce algorithm to calculate kinship coefficients to ensure similar degrees of relatedness among females^25,26^.

The baboons were raised and maintained on a standard monkey chow diet (high complex carbohydrates; low fat (“Monkey Diet 15%/5LEO,” LabDiet, PMI Nutrition International) prior to study initiation. Animals were acclimated together in the study cohort in a group cage for at least 8 weeks prior to study. Animals had free access to chow *ad libitum* in individual cages during feeding times. Food consumption, Na^+^ intake, body weight, menses cycle, and health status were monitored weekly for each animal throughout the study.

### Clinical measures related to kidney function and BP

A complete blood count with differential was performed at the start of the study. Baseline blood chemistry panels measured glucose, blood urea nitrogen (BUN), creatinine, total protein, albumin, globulin, cholesterol, ALT/SGPT, AST/SGOT, alkaline phosphatase, Na^+^, K^+^, chloride, carbon dioxide, and anion gap as described in Spradling-Reeves *et al*^18^. Serum 17 beta-estradiol was measured in female baboons with the 17 beta Estradiol ELISA Kit (Abcam); human female serum (age 28 years) was used as a positive control.

### Telemetry implantation, kidney sample, and blood collection

Animals were preoperatively treated with ketorolac (5-30 mg) preoperatively, sedated with ketamine (10mg/kg, IM), and maintained on isoflurane (1.3-3.0%) anesthesia throughout telemeter implant, kidney biopsy, and blood collection procedures. Animals were monitored during collections and throughout recovery. Postoperative care included observation for swelling at the surgical sight and monitoring until animal was conscious. Buprenorphine (0.2 mg/kg, SQ) was administered for post-operative pain relief immediately once animals were awake. Buprenorphine (0.2 mg/kg, SQ) was also administered as needed 64-48 hours after surgery. Blood was drawn in EDTA tubes, and kidney cortex biopsies were collected via ultrasound guidance and flash frozen in liquid nitrogen. Animals were euthanized the day after final kidney cortex biopsy collection by IV administration of >100mg/kg of Fatal-Plus (Vortech Pharmaceuticals, LTD.). A catheter was placed in the cephalic vein and Fatal-Plus was delivered followed by saline. Confirmation of death was by auscultation of the heart, checking for lose of anal tone and pupils being fixed and dilated. Bilateral thoractomy was performed as a secondary method of euthanasia. Kidney cortex and medulla samples were obtained during necropsy and each sample was placed in mesh cassettes with 10% neutral buffered formalin prior to embedding tissues in paraffin for IHC.

### Diet, housing, and BP measurements in baboons

Females were run into individual cages for 4 hours (passing over an electronic weighing scale) for feeding ad libitum once a day; they were fed a low sodium (LS) diet (17.3 mmol Na^+^/500 g food consumed) for 6 weeks followed by a high sodium (HS) diet (210 mmol Na^+^/500 g food consumed) for 6 weeks. Female baboons were housed in an outdoor social group with one vasectomized male (not on study) to provide full social and physical activity. Females were surgically implanted with a PhysioTel Digital implant model M10 (Data Science International (DSI)) under the external abdominal oblique muscle, and the catheter was implanted in the femoral artery to obtain 24-hour continuous BP, heart rate, temperature, and gyroscopic activity measurements in Ponemah software v 6.51 (DSI).

### Clinical measures related to kidney function and BP

Serum 17 beta-estradiol was measured using a randomized plate layout with 3 replicates per sample in female baboons on the LS and HS diets with the 17 beta Estradiol ELISA Kit (Abcam); human female serum (age 28 years) was used as a positive control.

### Telemetry data analysis

Ponemah software was used for BP analysis with signal acquired at a logging rate of 10 seconds per minute. Data were reduced to 64-hour periods throughout the study for mean arterial BP (MAP), systolic BP, diastolic BP, and heart rate. The data were calculated for 64-hours continuously, as well as daytime and nighttime intervals within the 64-hours, as observed in activity measurements. Blood pressure counts were determined by analysis of ECG waveforms. Nighttime intervals were determined by reduction in activity indicating sleep and dip in BP compared to daytime. BP data reported here are based on the last days of recording on each diet before kidney cortex biopsy collection.

### RNA isolation, library preparation, and sequencing

RNA was isolated from flash frozen kidney biopsy samples using the Direct-zol RNA Miniprep Plus Kit (Zymo Research, R2051). Samples were homogenized in 600 μL of TRI reagent (Zymo Research, R2051) using a BeadBeater (Biospec Products) for 3 x 30 sec with RNA purification according to the Direct-zol RNA Miniprep Plus Kit instructions. RNA was quantified by Qubit RNA BR Assay Kit (Invitrogen, Q10210), and RNA integrity was determined with an RNA ScreenTape Kit on the TapeStation 2200 system (Agilent, 5067-5576, 5067-5578, 5067-5577). cDNA libraries were prepared using the KAPA Stranded mRNA-Seq Kit (Roche, (07962193001). Libraries were quantified using a KAPA Library Quantification Kit (Roche, 07960204001), and quality was determined with a DNA ScreenTape Kit on the TapeStation 2200 system (Agilent, 5067-5582, 5067-5583, 5067-5586). Paired end sequencing was done using a HiSeq 2500 (Illumina). All samples were randomized during this process.

### RNA-Seq data processing

LS and HS transcripts were processed together in Partek^®^ Flow^®^ (Partek^®^, Inc.) representing over 9,300,000 reads per sample, with minimum read length of 25 and minimum Phred 30. Over 90% of reads were aligned using HISAT2 to the olive baboon genome (papAnu2.0; March 2012), with coverage >3 and depth >9. Data were normalized by TMM+1.Transcripts were annotated to identify kidney-specific cell types as shown by Park *et al*^10^.

### Weighted gene co-expression network analysis

Transcripts were filtered for total counts <14 among all samples prior to filtering based on coefficient of variation for the top 20,000 transcripts for each diet prior to performing diet-specific WGCNA. The R package WGCNA was run with Pearson correlation as described by Langfelder and Horvath^27^. WGCNA was selected as the most appropriate method for transcriptome analysis due to the ability to capture variation in gene expression data and identify clusters of interconnected transcripts represented by modules. Each module is then compared to a trait by correlation resulting in a correlation value and p-value which accounts for the average transcript significance across the module. This provides more information than simply binning two groups, as well as maintaining power and reducing the multiple testing burden. WGCNA modules were evaluated for validity by identification of known genes involved in biological processes related to traits they correlated with according to the GWAS catalog.

### Pathway and network analysis

Correlation directionality was used as a proxy for differential gene expression in pathway analysis and calculated by subtracting the values for each transcript of the animal with the lowest BP from the animal with the highest BP for significant modules. Pathway and causal network enrichment analyses using terms “salt-sensitive hypertension” for genes in significant WGCNA modules were performed using Ingenuity Pathway Analysis software (IPA; Ingenuity^®^ Systems) to identify direct connections with p-value cutoff <0.05.

Significant WGCNA modules, upstream regulators and master regulators identified from IPA were annotated with GWAS catalog variants associated with terms “hypertension” and “blood pressure” from the GWAS Catalog (Galaxy Genome, https://usegalaxy.org/) to identify genes with variants previously associated with BP.

### IHC

Kidney cortex and medulla tissues were obtained after necropsy. Tissues were placed in mesh cassettes and dropped into 10 volumes of 10% neutral buffered formalin and tissues were embedded in paraffin. Each paraffin block was cut to generate slides for IHC and one slide from each antibody was counterstained. All antibodies were obtained from Santa Cruz Biotechnology and used at a 1:500 dilution. Primary antibodies included Anti-PPARγ Antibody (E-8) (sc-7273), Anti-ACSVL4 Antibody (H-6) (sc-393309), Anti-Myc/c-Myc Antibody (9E10) (sc-40), Anti-VEGF Antibody (C-1) (sc-7269), Anti-CTL2 Antibody (F7) (sc-101266). All primary antibodies utilized a biotinylated secondary antibody from Vector Laboratories (Burlingame, CA. horse antimouse, catalog no. BA-2000, 1:1000 dilution). Standard IHC avidin-biotin-peroxidase complex technique (Elite ABC kits, cat.# PK-6100, Vector Labs: Burlingame, CA) was used to visualize protein expression using 0.02% DAB (3,3’-diaminobenzidine tetrahydrochloride) with 2.5% nickel sulfate as chromagen. Tissues were incubated in 60°C oven for 20 minutes, deparaffined with xylene 10 mins 3 times, and rehydrated twice in 100% EtOH for 5 mins. Tissues were microwaved in a. 0.01M citrate buffer (pH 6.0) to boil for antigen retrieval. Slides were positioned onto Sequenza racks and followed protocol as described with kit reagents. A total of 3 slides were used for IHC per antibody, and 6 images were captured of each slide. Images were obtained with a SPOT RT3 cooled color digital camera (2650 x 1920 pixels, Diagnostic Instruments, McHenry, IL, USA) mounted on a Nikon E600 microscope (Nikon, Inc., Melville, NY, USA). One slide from each protein was counterstained with H&E. Images were analyzed using ImageJ software with a background filter cutoff threshold of 200 to quantify staining presence and intensity. IHC staining demonstrated glomeruli or tubule specific protein expression, therefore each slide was normalized according to number of pixels containing glomeruli or tubules in the total image space. All samples were blinded upon staining and analysis.

### Statistics

#### Food consumption, Na^+^ intake, hormone, BP traits, and IHC

Mean values and standard deviations for food consumption and Na^+^ intake are reported as the average over 7 days prior to biopsy collection. Multiple unpaired t-tests with two-stage step-up Benjamini, Krieger, and Yekutieli method was performed on each BP traits^28^. Spearman correlation was performed on each of the food consumption, Na+ intake, hormone, and BP traits. IHC protein expression values were analyzed using Pearson correlations against the above traits. Data are expressed as mean, mean ± SD, or correlation r value, and were considered to be statistically significant if p-value < 0.05.

#### Transcripts

TMM+1 was used for normalization of trimmed and aligned transcripts. Multiple unpaired t-tests with two-stage step-up Benjamini, Krieger, and Yekutieli method were performed on cell type-specific transcripts between LS and HS samples^28^. WGCNA was performed using Pearson correlation with a p-value cutoff < 0.05. Correlation directionality was calculated by subtracting expression values across transcripts of animals with the lowest BP from animals with the highest BP for transcripts correlated with BP and input into as expression values for IPA network analysis. Upstream analysis and causal network regulator lists were ranked by p-value and target molecule number. Data are expressed as mean, mean ± SD, or correlation r value, and were considered to be statistically significant if p-value < 0.05.

## Results

### Food consumption, sodium intake, and serum hormone measurements

Healthy pedigreed females were identified from familial lines to identify a cohort with variation in BP in order to identify genetic variation correlated with BP variation (Supplementary Table S1, S2). Unlike male baboons, sodium sensitivity could not be predicted for sodium naïve females using sodium lithium counter transport measures (Supplementary Table S3)^18^. All baboons on study consumed similar amounts of food for the LS and HS diets (Figure 1A). As expected, Na^+^ intake was much greater on the HS diet than the LS diet (Figure 1B). Serum 17 beta-estradiol concentrations did not significantly differ between the diets (Figure 1C).

**Figure 1.**
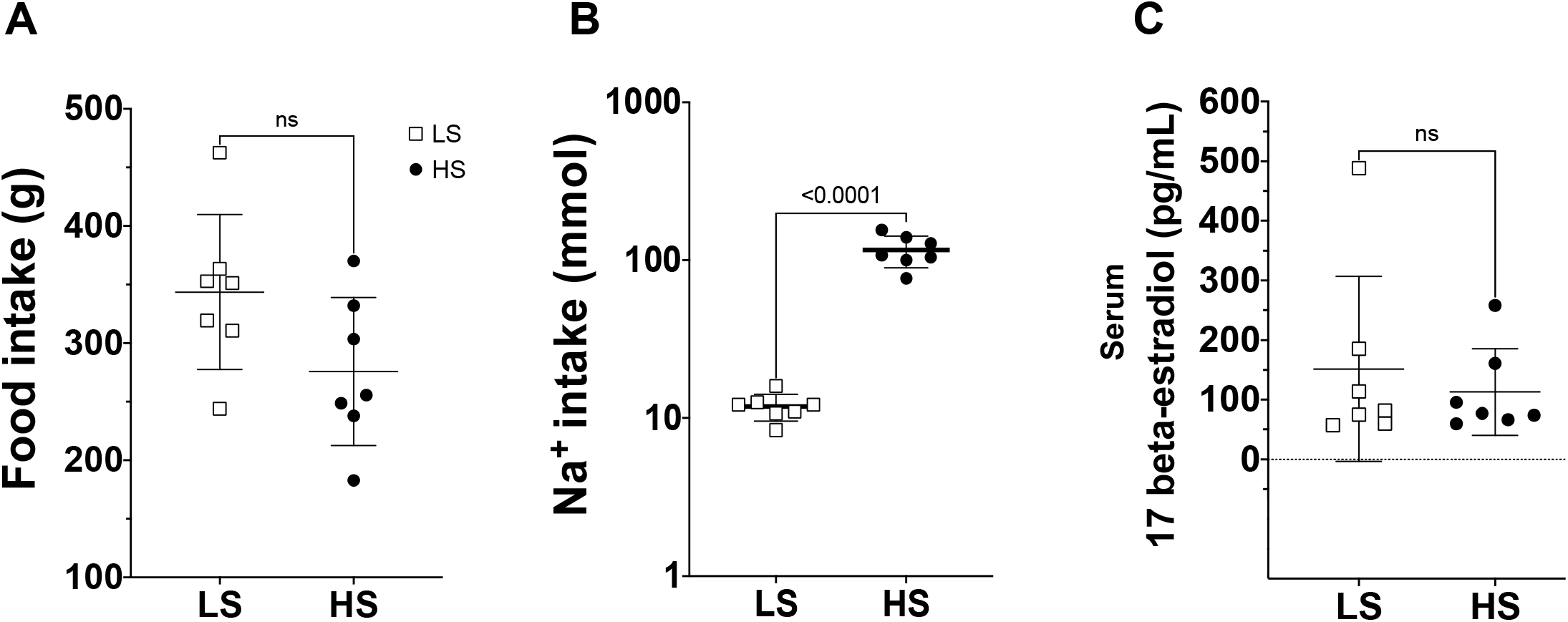
Diet and hormone measures on LS and HS diet. LS (square, n=7) and HS (circle, n=7) diets are shown for each measure on the x-axis. A. Food intake (g), B. Na^+^ intake (mmol), C. and 17 beta-estradiol (pg/mL) values for each animal are represented as points, and bars represent standard deviation. Significant differences between diets are labeled with p-value <0.0001 or “ns” if not significant.

### BP in sodium naïve female baboons with LS and HS diets

Telemetry measurements of MAP, diastolic BP, and systolic BP demonstrated BP is a continuous trait in this cohort for both diets, consistent with animal selection based on familial information. In addition, BP measures were significantly different among all animals in this cohort for the weekly 64-hour collections for both diets (Figure 2, Supplemental Table S4). The change in BP from LS to HS diet was significant for each animal across each measure (Figure 2A, 2B, 2C). Interestingly, some animals exhibited decreased BP, while others showed increased BP in response to the HS diet (Figure 2).

**Figure 2.**
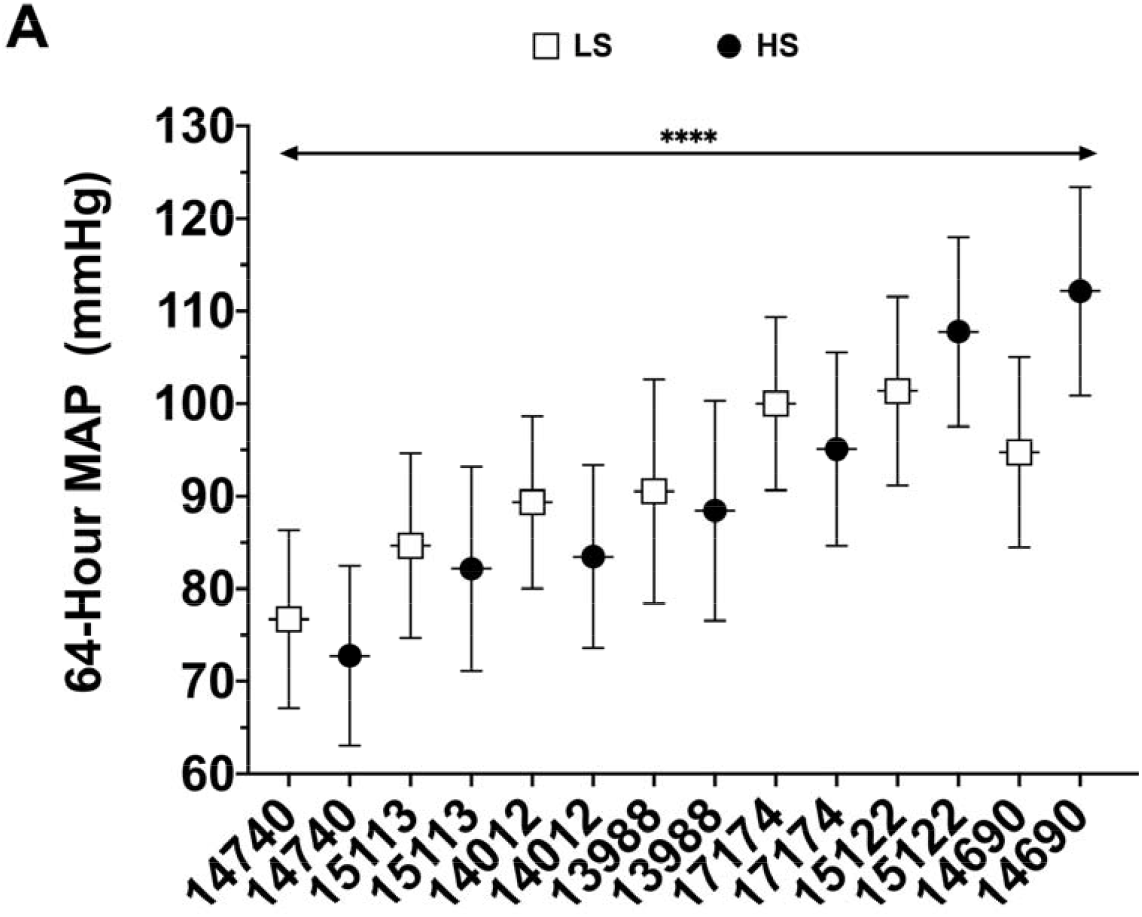
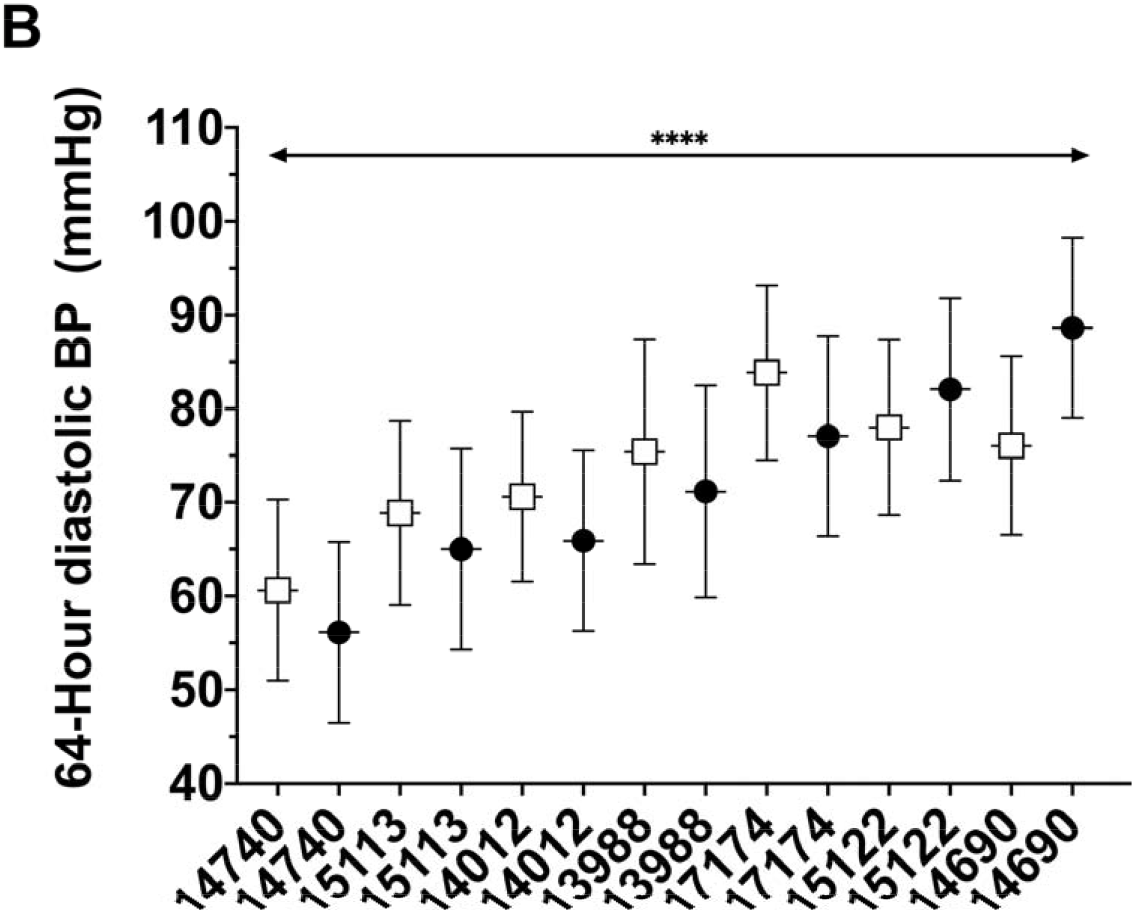
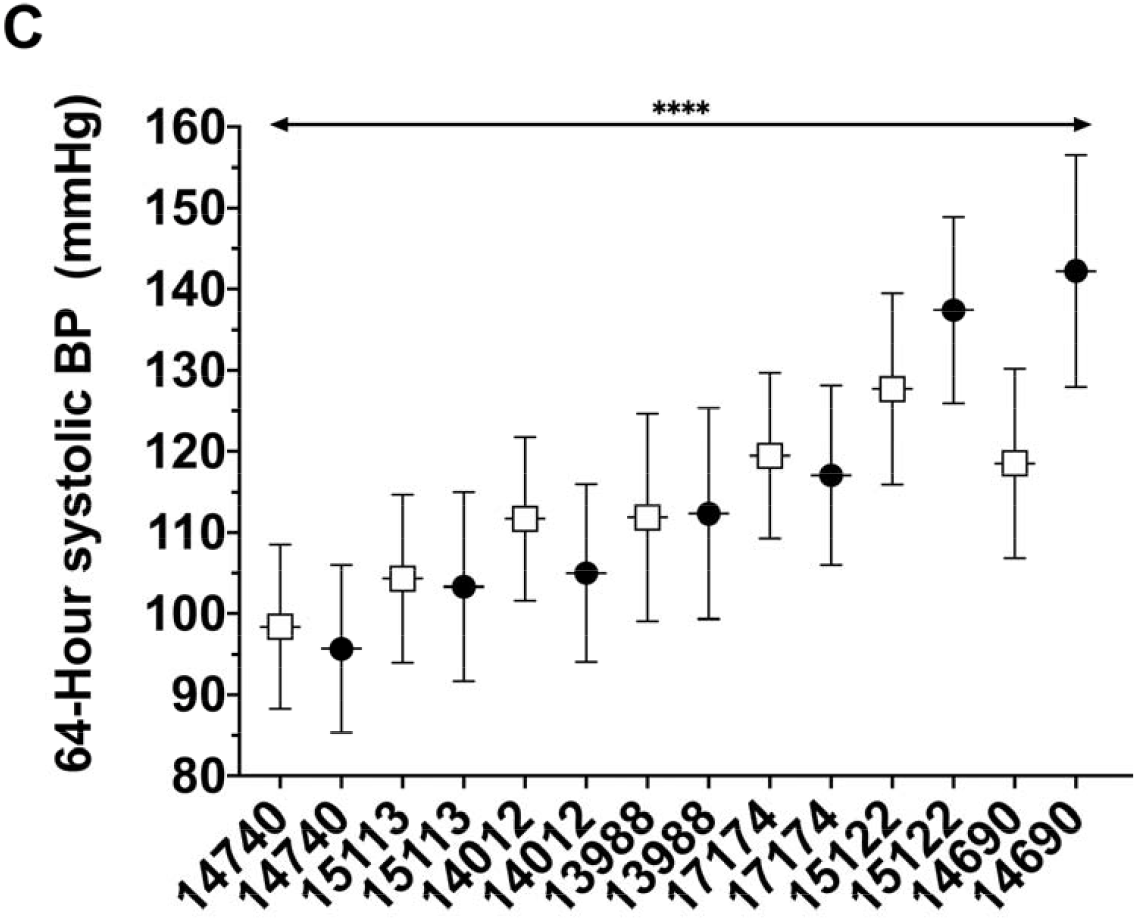
BP averaged over 64-hours each week for 6 weeks in female baboons on LS and HS diets. Animal ID is shown for each animal on the x-axis, and 64-hour BP (mmHg) is shown on the y-axis for A. MAP, B. diastolic BP, and C. systolic BP. Mean BP values represent at least 191,000 annotated BP counts for each animal, and are represented for LS as square and as a circle with standard deviation error bars. Significant differences in BP among all animals for both diets are indicated by **** (p-values <0.0001).

### Sodium and hormone correlations with BP in baboons

Na^+^ intake was significantly negatively correlated with serum 17 beta-estradiol concentrations (r=−0.77, p-value=0.0492) and with all BP measures on the LS diet (r < −0.86, p-values < 0.05) (Table 1, Supplementary Table). Serum 17 beta-estradiol concentrations were significantly positively correlated with 64-hour nighttime MAP, daytime systolic BP, and nighttime systolic BP for the LS diet (p-values=0.0238, 0.0480, 0.0238, respectively) (Table 1). No significant correlations were observed between BP and Na^+^ intake or serum 17 beta-estradiol concentration for the HS diet (Table 1).

**Table 1.** Correlations between Na+ intake, serum 17 beta-estradiol and BP in female baboons on the LS and HS diets with Spearman r, p-value, and sample size are shown.

|  |  | LS |  |  | HS |  |  |
| --- | --- | --- | --- | --- | --- | --- | --- |
| Correlation variables |  | r | p-value | N | r | p-value | N |
| Na<sup>+</sup> intake | 17 beta-estradiol | -0.77 | 0.049 | 7 | -0.71 | 0.088 | 7 |
| Na<sup>+</sup> intake | Nighttime MAP BP | -0.95 | 0.003 | 7 | -0.07 | 0.906 | 7 |
| Na<sup>+</sup> intake | Daytime systolic BP | -0.92 | 0.006 | 7 | -0.07 | 0.906 | 7 |
| Na<sup>+</sup> intake | Nighttime systolic BP | -0.95 | 0.003 | 7 | -0.07 | 0.906 | 7 |
| 17 beta-estradiol | Nighttime MAP BP | 0.86 | 0.024 | 7 | 0.64 | 0.139 | 7 |
| 17 beta-estradiol | Daytime systolic BP | 0.79 | 0.048 | 7 | 0.64 | 0.139 | 7 |
| 17 beta-estradiol | Nighttime systolic BP | 0.86 | 0.024 | 7 | 0.64 | 0.139 | 7 |

### Dietary differences in kidney cortex transcriptome

Kidney cortex RNA-Seq data showed 53,119 transcripts that passed quality filters for all samples from both diets. Merging our transcript list with kidney-specific transcripts, we found representation of all kidney cell types based on presence of transcripts unique to a single kidney cell type. Abundance of the most highly expressed kidney-specific genes (n=50) showed consistency in cell types represented from our ultrasound guided biopsy collections among study animals for both diets (Figure 3). Transcriptome profiles overall from LS diet and HS diet demonstrated differences in both gene expression extent of variation (*ANKRD1* coefficient of variation: LS, 2.30; HS, 0.075; *CR2* coefficient of variation: LS, 0.039; HS, 2.372) indicating there are significant transcriptome differences between the two diets.

**Figure 3.**
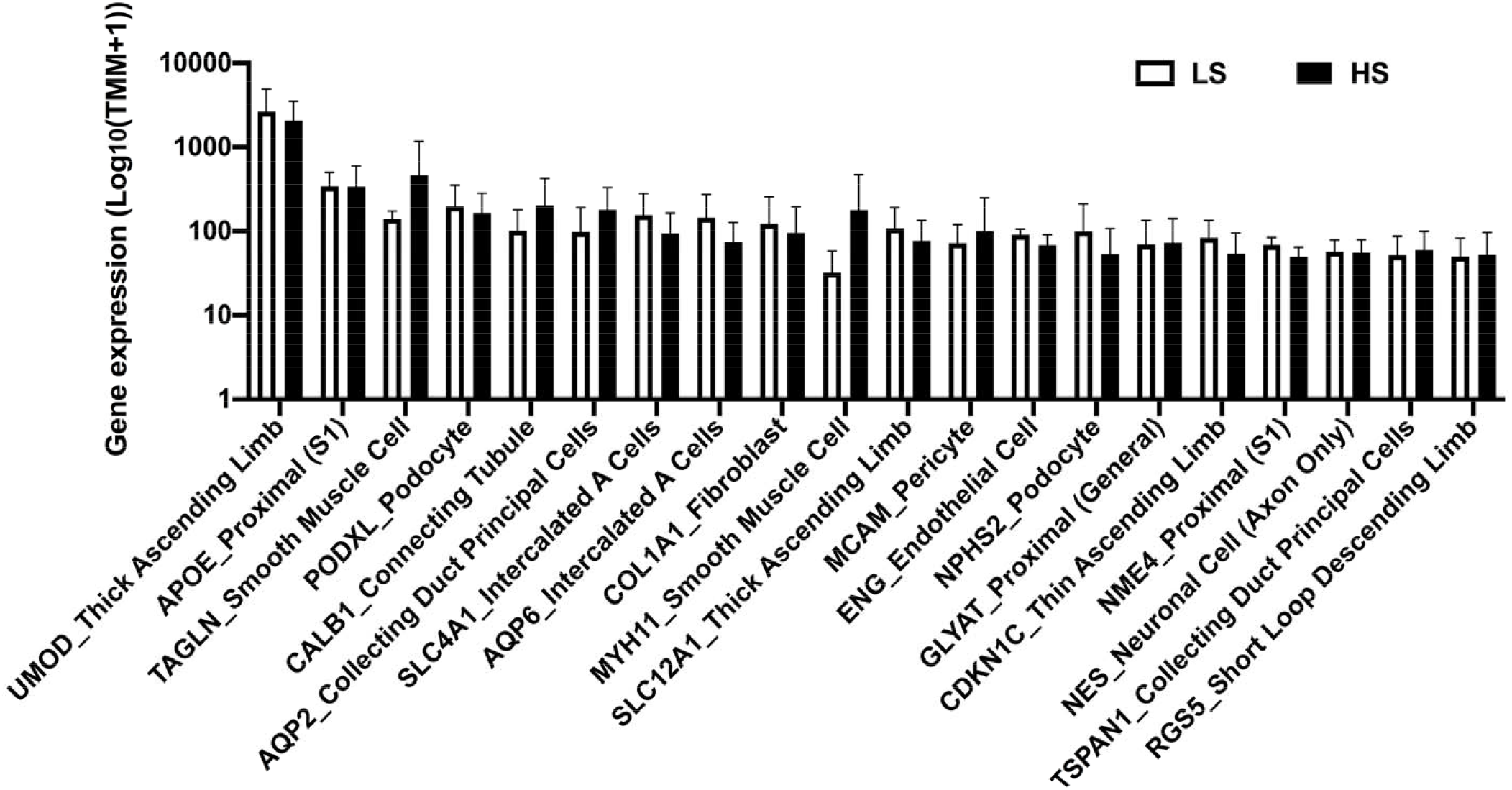
Cell type distribution of kidney-specific transcripts for LS and HS kidney cortex biopsies. Cell type distribution for each gene in LS (n=7, white columns) and HS (n=7, black columns) kidney cortex biopsies. Mean expression values were log(TMM+1) normalized and bars represent standard deviation. The x-axis is label includes gene ID and kidney cell type. No significant differences were observed between LS and HS diets for each gene by multiple unpaired t test.

### Correlations of kidney transcriptome with BP, sodium intake, and serum 17 beta-estradiol

WGCNA produced 13 modules of transcripts co-correlated with each other on the LS diet and 16 modules on the HS diet. Modules were subsequently tested for correlation with BP, Na^+^ intake, Na^+^ intake/animal weight, and serum 17 beta-estradiol measurements (Figure 4, Supplementary Figure S1, Supplementary Table 5).

**Figure 4.**
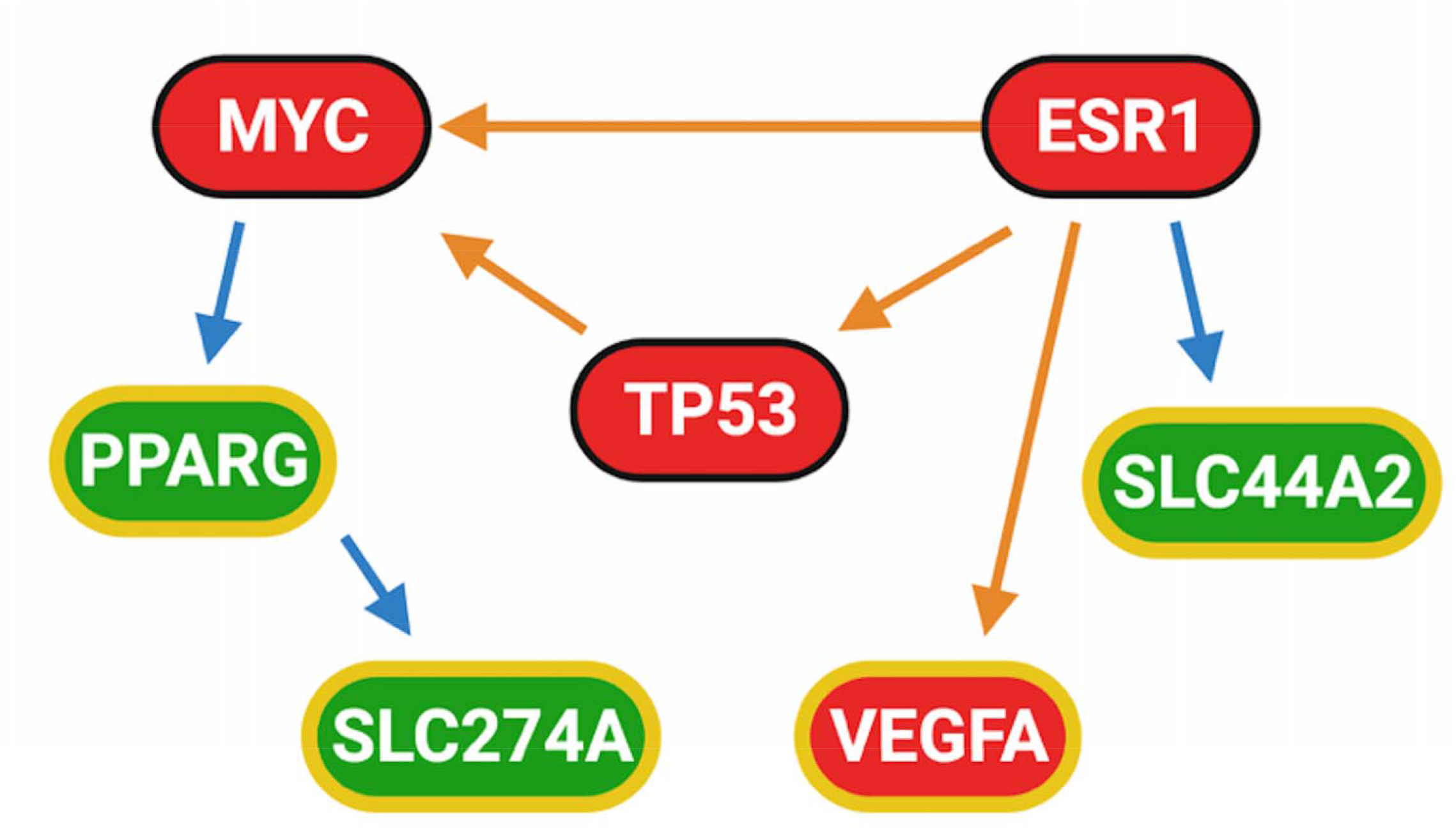
WGCNA of traits correlated with transcript modules in female baboons on HS diet (n=7). Each colored block in the left column represents a module of transcripts correlated with each other. The top values in each block indicate the correlation and the bottom values in each block indicate the p-value for each transcript module with the trait. The color scale to the right of the heatmap corresponds to correlation r values of each square. Abbreviated measures on the x-axis: Na^+^ intake (mmol Na^+^), Na^+^ intake/animal weight (mmol Na^+^/kg animal weight), 17 beta-estradiol (pg/mL), daytime MAP (mmHg), nighttime MAP (mmHg), 64-hour MAP (mmHg), daytime systolic BP (mmHg), nighttime systolic BP (mmHg), 64-hour systolic BP (mmHg), daytime diastolic BP (mmHg), nighttime diastolic BP (mmHg), and 64-hour diastolic BP (mmHg).

For the LS diet, we found one module correlated with Na^+^ intake/animal weight (orange n=58 transcripts, p-value=0.008) and one module correlated with 17 beta-estradiol (yellow, n=2283 transcripts, p-value=0.002); however, no modules were significantly correlated with BP measures (Supplementary Figure S1). For the HS diet, we found and one module correlated with Na^+^ intake/animal weight (pink, n=3064 transcripts, p-value=0.007) and one module correlated with 17 beta-estradiol (green, n=1347 transcripts, p-value=0.02), but this neither of these modules correlated with Na^+^ intake or BP. In addition, we found two modules (cyan, n=1166 transcripts; dark green, n=324 transcripts) positively correlated with BP traits: nighttime MAP, 64-hour diastolic BP, nighttime diastolic BP; nighttime diastolic BP; 64-hour systolic BP, and nighttime systolic BP (p-values=0.02, 0.05, 0.01; 0.04; 0.04, 0.03, 0.03, respectively) (Figure 4). These modules contained 183 renal-specific genes and 767 genes were annotated with 5733 GWAS variants associated with BP (Supplementary Table S5). Neither of these BP-related modules correlated with any clinical measures for the HS diet.

### Network analysis of kidney cortex transcripts correlated with BP on HS diet

Network analysis of modules correlated with BP for the HS diet (cyan, dark green) showed that the significant causal networks with the greatest number of direct downstream targets included genes related to hormone receptors (*ESRRG*, *ESRRA*, *ESR1*, *PGR*, *AR*), proliferation and differentiation (*TP53*, *MYC*, *ID2*, *GATA2*, *EP300*, *MITF*, *RB1*), methylation (*KDM4B*, *EZH2*), hypoxia (*HIF1A*), insulin and lipid regulation (*TCF7L2*, *SREBF1*), and inflammation (*PPARGC1*, *PPARA*, *PPARG*, *PPARD*, *PPARGC1B*, *NR4A1*) (Figure 5, Supplementary Figure S2, Supplementary Table S6, S15). Many targets of these regulators were related to sodium and ion management in the cell membrane, vascularization, and renin homeostasis. Notably 11 solute carriers including *AQP2*, 5 solute channels including *SCNN1A*, 2 ATPases, *RENBP*, and *VEGFB* were all down regulated downstream of the master regulators (Supplementary Table S7). In addition to these specific genes known to be related to blood pressure regulation, the vast majority of genes in our dataset showed an inverse correlation with BP on the HS diet (Supplementary Table S5). From the kidney cell specific expression data, transcripts annotated as inner medullary collecting duct cells were predicted to play important regulatory roles in upstream regulators correlated with BP variation. *ESR1*, *PAX2*, *CASZ1*, *MMT3*, and *MEF2B*, *EBF1*, *HDAC7*, *WWP2*, *PKN2*, *TCF7L2*, *NRIP1*, *PTEN*, *NCOR2*, *PLCG1*, *SETD7*, *EHMT2* were among the list of master regulators that correlated with BP and previously have been associated with BP from GWAS studies (Supplementary Table S6, S7).

**Figure 5.**
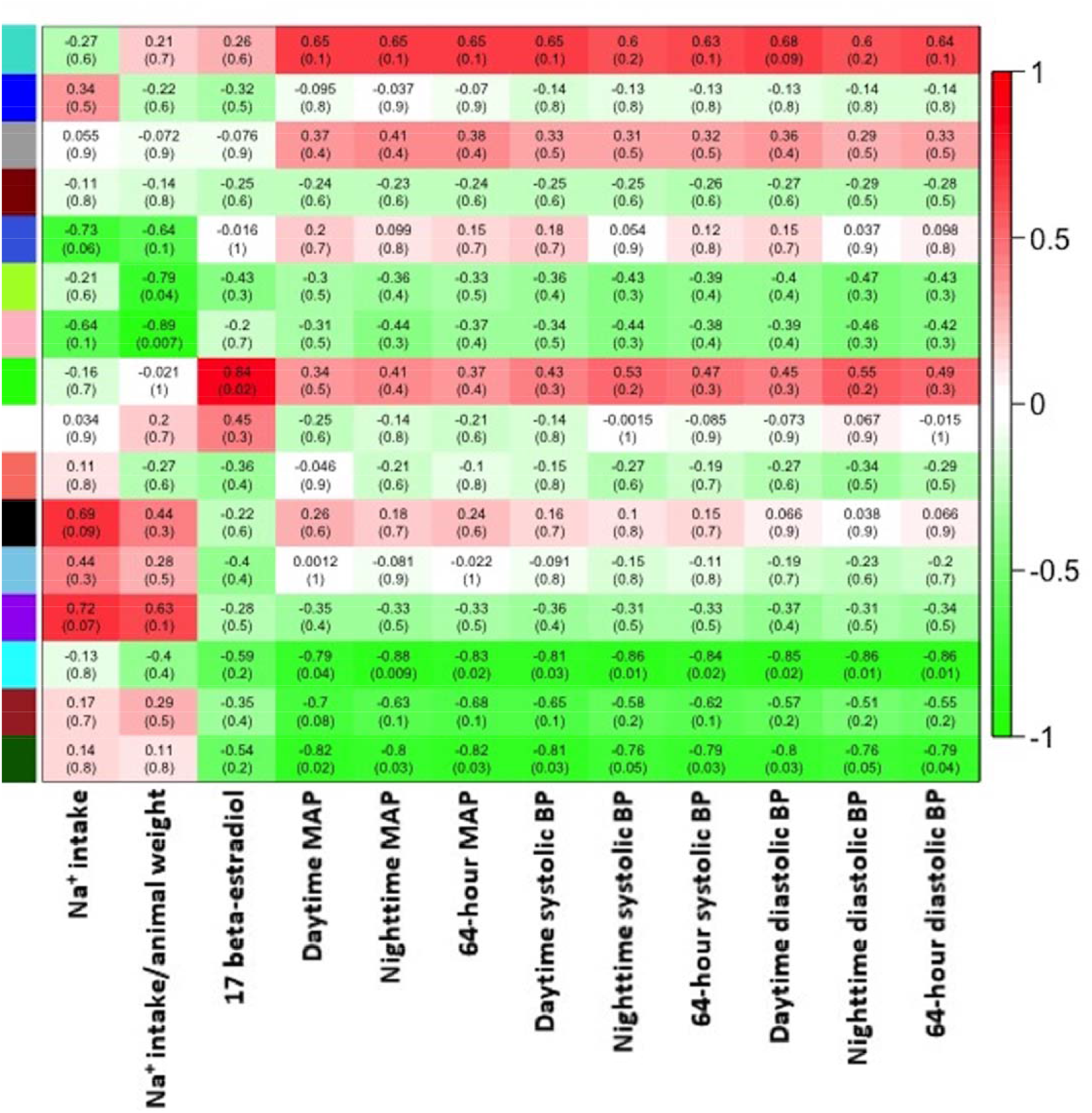
Network analysis of WGCNA transcripts correlated with blood pressure on HS diet. Upstream regulators of transcripts and targets are shown. Red shapes are genes predicted as activators and are positively correlated with blood pressure in the dataset. Green shapes are genes predicted as inhibitors, and genes in green are negatively correlated with blood pressure in the dataset. Arrows indicate direction of activation of downstream gene, and T lines represent inhibition of downstream gene. Orange lines indicate target activation supported by the literature, blue lines indicate target inhibition supported by the literature. Shapes outlined in yellow indicate genes selected for protein quantification by IHC.

### Immunohistochemistry of predicted network regulators

Protein expression from IHC of transcripts identified in network analysis (PPARG, ACSVL4, C-MYC, CTL2) in kidney cortex after HS diet demonstrated a distinct pattern differing between glomeruli and tubules (Figure 6, Supplementary Figure 3). IHC of tubules in the kidney medulla also exhibited expression of the same targets as the kidney cortex and were quantified for correlation analysis. Pearson correlation revealed a positive significant correlation between Na^+^ intake and ACLVS4 in cortex as well as a negative correlation between Na^+^ intake VEGFA in cortex tubules (p-values=0.048, 0.048, respectively). VEGFA in cortex tubules also significantly correlated with 17 beta-estradiol (p-value= 0.007) (Table 2). VEGFA in cortex glomeruli significantly correlated negatively with SLC274A transcript (p-value= 0.024). VEGFA in medulla correlated with C-MYC cortex (0.024). Pearson correlation of both ACLVS4 and CTL2 demonstrated significant positive correlation between protein expression in the cortex and medulla (p-value=0.019, 0.031). VEGFA in both cortex tubules and glomeruli was also positively correlated with 17 beta-estradiol (p-value= 0.006, 0.008). C-MYC in medulla correlated with PPARG in the Medulla (p-value=0.010). VEGFA cortex glomeruli also correlated positively with nighttime MAP BP and nighttime systolic BP (p-value=0.039, 0.024). None of the proteins measured in the kidney cortex or medulla correlated the gene expression of the encoding gene within networks correlated with BP identified by WGCNA (Supplementary Figure S2).

**Figure 6.**
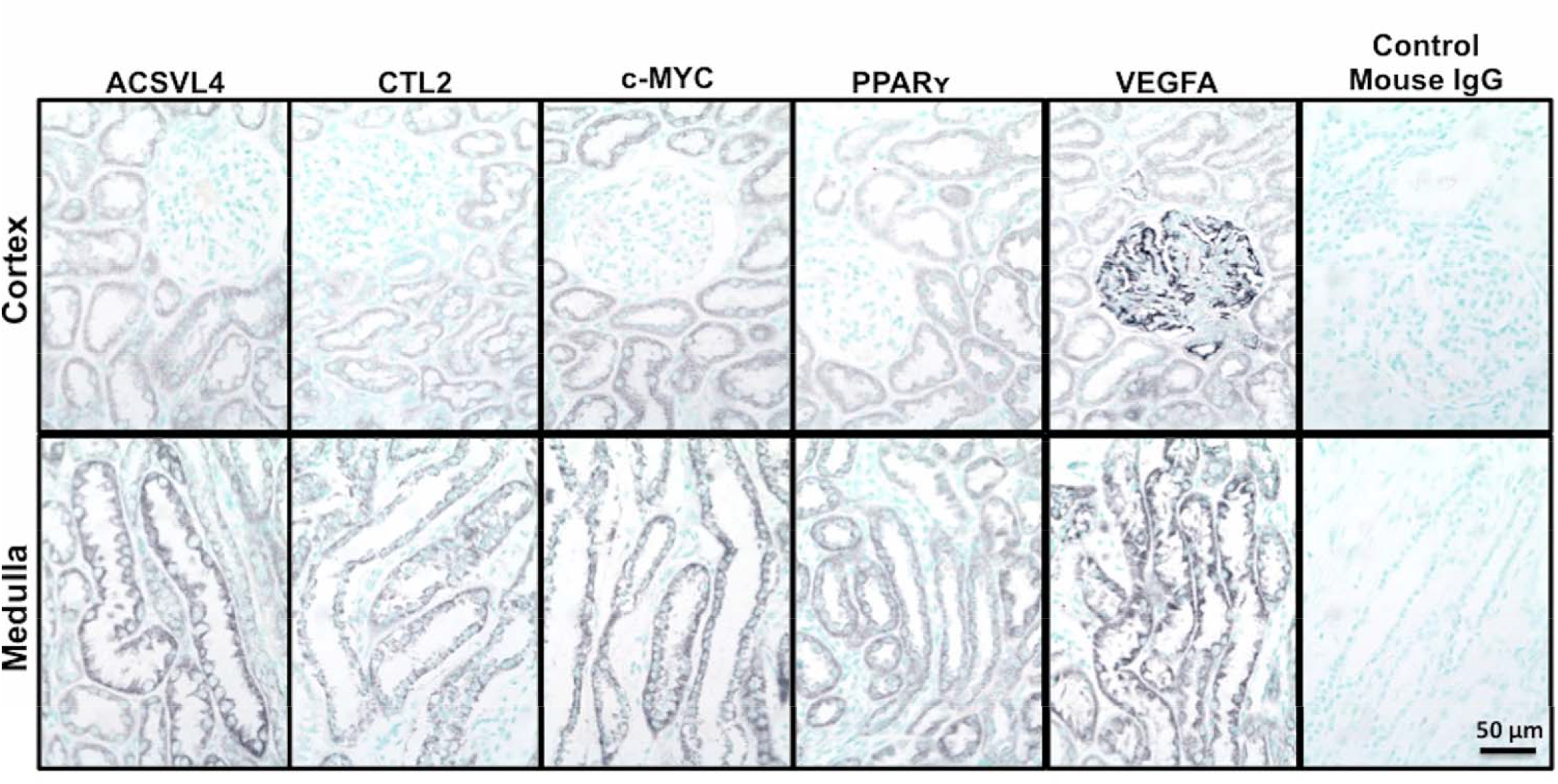
Protein expression of targets from network analysis in kidney cortex and medulla of female baboons on a HS diet. Representative images of kidney cortex and medulla sections stained for PPARy, ACSVL4, c-MYC, CTL2, and Control Mouse IgG. In each panel the grey area demonstrates protein expression, while the blue is methyl green stained background, not representative of protein expression. Staining demonstrated distinct patterns of expression observed in tubules and glomeruli for each target, when compared with control.

**Table 2.** Correlations between IHC values and BP related traits in female baboons on a HS diet. Correlation variables compared for each correlation with Pearson r and p-value are shown, for n=7 animals.

| Correlation variables |  | Pearson r | p-value |
| --- | --- | --- | --- |
| VEGFA Cortex Tubules | Na<sup>+</sup> intake | -0.56 | 0.189 |
| VEGFA Cortex Tubules | 17 beta-estradiol | 0.89 | 0.006 |
| VEGFA Cortex Glomeruli | 17 beta-estradiol | 0.89 | 0.008 |
| VEGFA Cortex Glomeruli | Nighttime MAP BP | 0.78 | 0.039 |
| VEGFA Cortex Glomeruli | Nighttime systolic BP | 0.82 | 0.024 |
| VEGFA Cortex Glomeruli | SLC274A Transcript | -0.74 | 0.058 |
| VEGFA Medulla | C-MYC Cortex | -0.80 | 0.030 |
| Na<sup>+</sup> intake | ACLVS4 Cortex | 0.67 | 0.101 |
| ACLVS4 Cortex | ACLVS4 Medulla | 0.84 | 0.019 |
| CTL2 Cortex | CTL2 Medulla | 0.80 | 0.031 |
| C-MYC Medulla | PPARG Medulla | 0.87 | 0.010 |
| PPARG Cortex | Menstrual Cycle | 0.93 | 0.002 |

## Discussion

A HS diet is known to influence BP and HT risk^1,2^. Previous studies in primates to identify molecular networks dysregulated by HS diet focused on males. Current clinical guidelines do not offer sex-specific treatment plans for sodium sensitive HT. The goal of this study was to understand the impact of HS intake on gene networks in the kidney that correlate with BP in female primates with the long-term goal of translating findings to women. To address this, sodium naïve female baboons were challenged with a HS diet, BP quantified by telemetry, and the renal transcriptome quantified by RNA-Seq in a cohort of baboons with significant variation in sodium sensitive BP. This study, which used the transcriptome as a tissue specific read-out to identify genetic variation relevant to kidney and BP response to HS diet, leveraged variation in BP as a first step to identify correlated kidney regulatory networks that may influence BP in female primates.

For the LS diet, sodium intake per kg of animal weight was similar to previous LS diets in humans. The amount of sodium in the HS diet used in this study falls within the range consumed by humans NHANES^29^. For the HS diet, sodium intake per kg of animal weight was comparable to a study of male baboons, approximately 2-3 times greater than the average sodium consumption in humans, but within the normal range of consumption in humans and about 16-fold less than HS diets used in rodent studies^18,30–32^. Our results, as well as study results in male baboons, demonstrate this amount of sodium for the study period is sufficient to induce a molecular and physiological response.

Interestingly, systolic, diastolic, and MAP BP increased in some animals and decreased in others when comparing HS diet BP with LS diet BP, similar to observations in the human population^2,33^. BP, Na^+^ intake, and serum 17 beta-estradiol correlated with each other on the LS diet, indicating a LS diet is not always beneficial for BP as has been observed in human studies^33^. Correlations of BP with 17 beta-estradiol were not seen on the HS diet, emphasizing the complex interplay of environmental factors such as dietary sodium and sex hormones on BP as shown in human populations ^18–21,30,34,35^.

Cell type composition of bulk RNA-Seq kidney cortex biopsy samples was assessed using abundance data from genes known to be expressed only in a single kidney cell-type^36^. This method increased our confidence in the consistency of the kidney cortex biopsy sampling in all animals for both diets. Evaluating sampling consistency was essential for ensuring variation observed in gene expression was biological variation rather than variation from biopsy collections.

As mentioned previously, animals were not selected for “high” and “low” BP, but rather selected to capture variation in BP, reflecting BP as a continuous trait. Comparing HS diet BP to LS diet BP, some animals increased BP and some decreased BP. To leverage this variation, WGCNA was utilized as an unbiased approach to identify kidney transcripts in which variation correlated with variation in BP for both diets. The kidney transcriptome revealed modules of genes correlated with BP measures including nighttime MAP, 64-hour diastolic BP, nighttime diastolic BP; nighttime diastolic BP; 64-hour systolic BP, and nighttime systolic BP only for the HS challenge.

Network analysis of the highest BP animals versus the lowest BP animals on the HS diet showed activation of processes related to inflammation and fibrosis, and decreased ion homeostasis, vascular remodeling, vascular flexibility, hypoxic response, sex hormones, and lipid metabolism. For example, the network included *EZH2*, *a* pro-fibrotic and anti-angiogenic gene^37^ and included estrogens, androgens, and their receptors, which have been shown to play a roles in renal ion homeostasis affecting the RAAS^38,39^. Furthermore, network genes *TP53*, *MYC*, *ID2*, *RB1*, *GATA2*, *EP300*, *MITF*, *CTNNB1*, and *SNAI1* play roles in proliferation, remodeling, and fibrosis, potentially leading to less vascular flexibility^40^. The network also contained HIF1A, a regulator of hypoxic response, which is inhibited by a HS diet in rats^41^. Reduction of O_2_ inhibits the Na-K-ATPase complex, which is responsible for Na+ reabsorption and regulation of lipids and glucose^42^. Buildup of fatty acids is also shown to lead to acute kidney injury, triggering fibrosis^43^. Our findings suggest that animals with increased BP with the HS diet have decreased Na^+^ resorption, and with prolonged HS diet will develop kidney injury and fibrosis. Our findings of the PPAR family inclusion in our network further supports these findings with their role in lipid regulation mediated by hormones, which lead to increased in inflammation when down regulated^43^. Taken together, our findings indicate a MYC and ESR1 regulated network, that influences Na^+^ ion absorption, inflammation, and fibrosis, is central to the kidney response in animals with increased BP compared to animals with decreased BP.

To determine protein abundance and cell localization of key molecules in the regulatory network, we performed IHC on kidney samples collected at the end of the HS diet challenge. IHC showed different distributions of proteins within glomeruli and tubules. Of greatest interest was the downstream target VEGFA which had been associated with systemic sodium sensitivity, and predicted upstream regulator MYC. VEGFA protein abundance correlated with BP measures and plasma 17 beta-estradiol concentrations ^44^. While c-MYC did not correlate with BP, it did correlate with VEGFA abundance. Although gene expression is often used as a proxy for protein activity, many key regulators’ activity-such as MYC-are dependent on post translational modifications ^45,46^. Consequently, lack of protein abundance correlation with BP variation does not contradict the putative roles of these regulators in the network. These findings further support our network and previous studies in which an activation cascade of upstream regulators beginning with ESR1 activation by estrogen leads to upregulation of MYC and TP53^44,47–49^. MYC is known to then bind directly to the VEGFA promoter, and lead to increased angiogenesis^49^. c-MYC in the kidney cortex negatively correlated with VEGFA in the medulla, suggesting that the relationship between *MYC* and *VEGFA* may not be kidney cortex specific. ESR1 has also been shown to directly regulate VEGFA in other tissues^50^. Furthermore, these relationships were kidney cell specific, including the cortex where specific substructural relationships in the glomeruli and tubules were observed.

VEGFA is also of interest having previously been demonstrated to impact BP. Patients with VEGFA inhibition typically develop sodium dependent HT which is thought to be due in part to impaired sodium excretion^51–53^. Previous studies in normotensive Wistar–Kyoto rats also demonstrated a relationship between VEGFA inhibition and a HS diet on BP^54^. Rodents with VEGFA inhibited while on a HS diet exhibited the highest BP (15 mmHg higher) compared to controls^53^. While many studies have shown that multiple VEGF family members may contribute to lowering BP, it is often difficult to tease out the role specific family members ^55,56^. While the association between VEGF and BP is not novel, the putative role of increased VEGFA in the kidney cortex and increased BP is novel. This observation indicates the female primate kidney response to sodium differs from the systemic relationship between BP and VEGFA previously observed in humans and rodents.

Our study shows a relationship between BP, sodium diet, and estrogen in female primate kidneys for a LS and a HS diet. We identified ESR1 and MYC as novel putative regulators of BP and provided evidence suggesting that VEGFA is an important player in BP regulation on a HS diet in female primate kidneys. Overall, these results provide a first step towards validating the *in silico* networks identified from the transcript data that are predicted to regulate BP response to a HS diet. Further validation includes identification of genetic variants in the genes composing this central regulatory network, and validation of the mechanistic role of these novel putative regulators of BP, which may provide therapeutic targets for HT prevention and treatment.

## Availability of data and materials

The datasets generated and/or analyzed during the current study are available in the locations detailed below:

RNA-Seq dataset: NCBI GEO GSE181248; https://www.ncbi.nlm.nih.gov/geo/

GWAS data were derived from: GWAS Catalog; https://www.ebi.ac.uk/gwas/

Cell type specific gene identifiers were derived from Park *et al*^10^.

## Acknowledgements

None

## Additional Files

Supplemental file names:

Data Supplement 1.xls

Data Supplement 2.docx

