## Supplemental Figures 1-3 for "Blood pressure and the kidney cortex transcriptome response to high sodium diet challenge in female nonhuman primates"

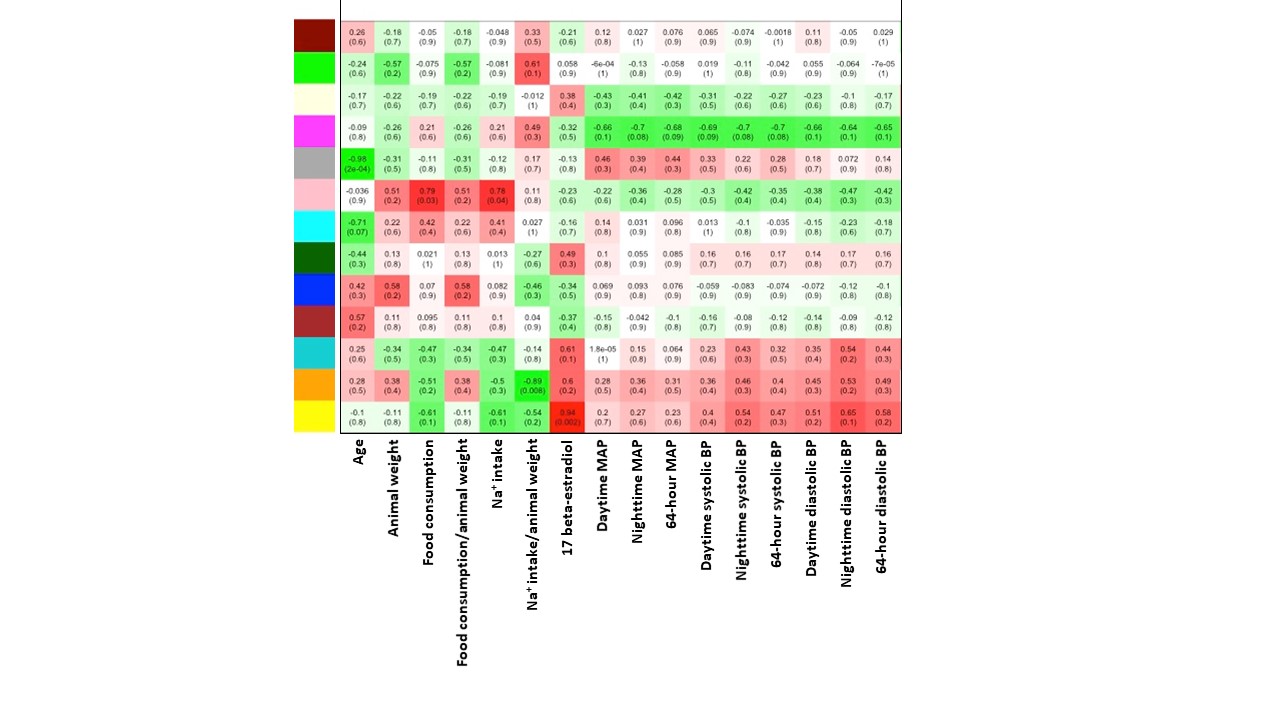

**Supplementary Figure S1. WGCNA of traits correlated with transcript modules in female baboons on LS diet (n=7). Each colored block in the left column represents a module of transcripts correlated with each other. The top values in each block indicate the correlation and the bottom values in each block indicate the p-value for each transcript module with the trait. The color scale to the right of the heatmap corresponds to correlation r values of each square. Abbreviated measures on the x-axis: Age, Animal weight, Na^+^ intake (mmol Na^+^), Na^+^ intake/animal weight (mmol Na^+^/kg animal weight), 17 beta-estradiol (pg/mL), daytime MAP (mmHg), nighttime MAP (mmHg), 64-hour MAP (mmHg), daytime systolic BP (mmHg), nighttime systolic BP (mmHg), 64-hour systolic BP (mmHg), daytime diastolic BP (mmHg), nighttime diastolic BP (mmHg), and 64-hour diastolic BP (mmHg).**

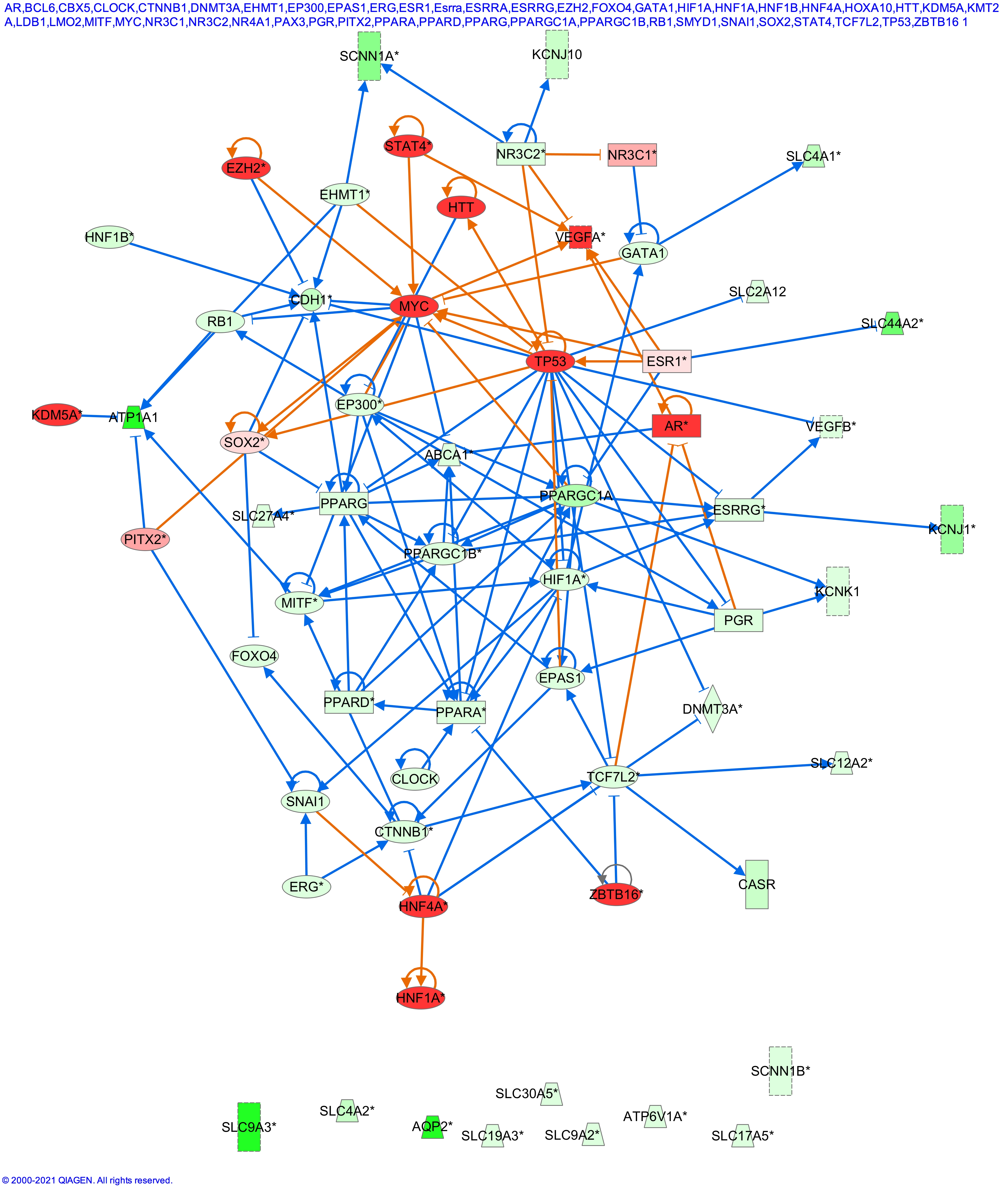

**Supplementary Figure S2. Network analysis of WGCNA transcripts correlated with blood pressure on HS diet. Upstream regulators of transcripts and targets are shown. Molecules in orange are genes predicted as activators and are positively correlated with blood pressure in the dataset. Molecules in blue are genes predicted as inhibitors, and genes in green are negatively correlated with blood pressure in the dataset. Arrows indicate direction of activation of downstream gene, and T lines represent inhibition of downstream gene. Orange lines indicate target activation supported by the literature, blue lines indicate target inhibition supported by the literature.**

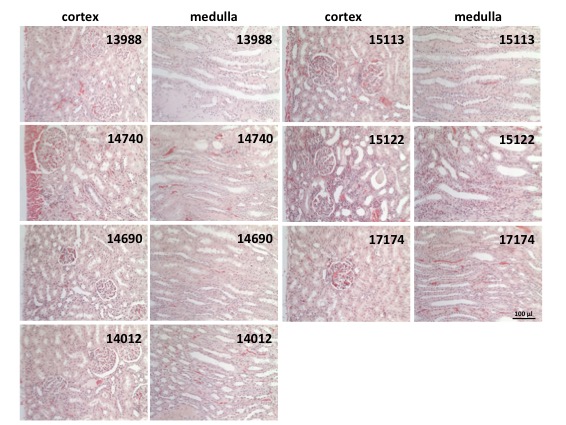

**Supplementary Figure S3. H&E staining of kidney cortex and medulla in female baboons on a HS diet.** Representative images of kidney cortex and medulla sections stained with H&E. Staining demonstrated distinct patterns of tubules and glomeruli in kidney cortex samples for each animal. Kidney medulla sections only exhibited tubules in each animal.
